# Goodness of fit checks for binomial N-mixture models

**DOI:** 10.1101/194340

**Authors:** Jonas Knape, Debora Arlt, Frédéric Barraquand, Åke Berg, Mathieu Chevalier, Tomas Pärt, Alejandro Ruete, Michał Zmihorski

## Abstract

1. Binomial N-mixture models are commonly applied to analyze population survey data. By estimating detection probabilities, N-mixture models aim at extracting information about abundances in terms of actual and not just relative numbers. This separation of detection probability and abundance relies on parametric assumptions about the distribution of individuals among sites and of detections of individuals among repeat visits to sites. Current methods for checking assumptions are limited, and their computational complexity have hindered evaluations of their performances.
2. We develop computationally efficient graphical goodness of fit checks and measures of overdispersion for binomial N-mixture models. These checks are illustrated in a case study, and evaluated in simulations under two scenarios. The two scenarios assume overdispersion in the abundance distribution via a negative binomial distribution or in the detection probability via a beta-binomial distribution. We evaluate the ability of the checks to detect lack of fit, and how lack of fit affects estimates of abundances.
3. The simulations show that if the parametric assumptions are incorrect there can be severe biases in estimated abundances: negatively if there is overdispersion in abundance relative to the fitted model and positively if there is overdispersion in detection. Our goodness of fit checks performed well in detecting lack of fit when the abundance distribution is overdispersed, but struggled to detect lack of fit when detections were overdispersed. We show that the inability to detect lack of fit due to overdispersed detection is caused by a fundamental similarity between N-mixture models with beta-binomial detections and N-mixture models with negative binomial abundances.
4. The strong biases in estimated abundances that can occur in the binomial N-mixture model when the distribution of individuals among sites, or the detection model, is mis-specified implies that checking goodness of fit is essential for sound inference in ecological studies that use these methods. To check the assumptions we provide computationally efficient goodness of fit checks that are available in an R-package nmixgof. However, even when a binomial N-mixture model appears to fit the data well, estimates are not robust in the presence of overdispersion unless additional information about detection is collected.

## 1 Introduction

Count surveys are often conducted as parts of population monitoring programs and ecological studies to follow changes in abundance of organisms in the wild. N-mixture models (Royle 2004; Royle & Dorazio 2006) have become increasingly applied to data from count surveys to correct for imperfect detection and yield estimates of absolute abundances instead of just relative abundances. These models are intuitively appealing because they can be applied to data from surveys with simple as well as more complex field protocols and allow simultaneous inclusion of explanatory variables for both abundance and detection processes.

N-mixture models are hierarchical models composed of two layers where the first layer gives a statistical model for the distribution of individuals among sampled sites and the second layer a statistical model for the sampling or detection process. Binomial N-mixture models (Royle 2004) are a particular class of models that rely only on repeated counts from a large number of sites to estimate absolute abundance while accounting for imperfect detection using binomial detection models. The assumptions of these models include that the population size is the same at each repeat visit to the same site, usually called the closure assumption, and that each individual could potentially be detected at each visit; that the distribution of the number of animals at each site is randomly and independently distributed according to some parametric distribution; and that all individuals are detected independently. The simplicity of collecting data under the protocol of the binomial N-mixture model has led some authors to suggest monitoring programs to incorporate multiple visits to sites (Lyons *et al.* 2012), while others have advised careful scrutiny of model performance before adopting the binomial N-mixture model for inferences (Hunt *et al.* 2012; Couturier *et al.* 2013). In the remainder of the paper we will refer to binomial N-mixture models simply as ‘N-mixture models’.

Because N-mixture models rely on parametric distributions and other assumptions, it is vital for reliable inference to investigate how sensitive estimates are to deviations from assumptions, and to devise methods for checking any assumptions that the models are sensitive to. N-mixture models have been shown to be reasonably robust to individual heterogeneity in detection unless detection probabilities are small (Veech *et al.* 2016), but to be sensitive to the closure assumption with overestimation of abundance when the assumption is violated (Toribio *et al.* 2012). Martin *et al.* (2011) showed through simulation that abundance was severely overestimated with an N-mixture model when detection probabilities were varying randomly among visits according to a nearly uniform distribution. They associated such variation, causing overdispersion in detection relative to the binomial distribution, with correlated behaviour among animals. They suggested a beta-binomial detection model to deal with it.

Many applications of N-mixture models use a Poisson abundance mixture which leads to a restrictive variance-mean scaling such that the variances of counts as well as abundances are proportional to their respective means. However, overdispersion is a common feature of population count data (Hoef & Boveng 2007; Lindén & Mäntyniemi 2011) and Taylor’s power law, with empirical as well as theoretical support, suggests that the variance-mean scaling of abundances follows a power law with an exponent typically somewhere between 1 and 2 (Cohen *et al.* 2013). Other work has suggested that the abundance distributions found in population surveys can be highly complex and irregular, effectively defying parametric modelling altogether (Dorazio *et al.* 2008; Canale & Prünster 2017). Abundance overdispersion is sometimes incorporated in N-mixture models by assuming a zero inflated Poisson, negative binomial or Poisson log-normal abundance mixture. Several studies have found estimates from N-mixture models applied to survey data to depend on which abundance mixture is used (Kéry *et al.* 2005; Joseph *et al.* 2009) and that estimates from models (Royle 2004) with a negative binomial abundance mixture sometimes behave poorly, yielding infinite maximum likelihood estimates of abundance (Dennis *et al.* 2015). This has led to recommendations for using zero inflated Poisson mixtures instead of negative binomial mixtures, even if the latter provide a better model fit (Joseph *et al.* 2009; Kéry & Royle 2016). Seemingly more realistic estimates do however not necessarily translate into better inference as the use of an ill fitting model could result in misleading conclusions.

In relation to their common usage relatively few studies have examined the performance of N-mixture models (Dennis *et al.* 2015), and calls have been made for evaluating and developing methods for assessing their fit (Kéry & Royle 2016; Knape & Korner-Nievergelt 2016). Our aim in this paper is to propose a set of tools, including graphical checks and overdispersion measures, to assess goodness of fit of N-mixture models, and to evaluate their ability in detecting lack of fit when there is overdispersion in abundance or detection relative to the fitted model. The graphical checks are based on randomized quantile residuals (Dunn & Smyth 1996; Warton *et al.* 2016), which have recently been applied to check goodness of fit of occupancy models (Warton *et al.* 2017), while the overdispersion measures are defined through two types of chi-square statistics. Compared to previously suggested goodness of fit checks that require parametric bootstrapping (Kéry *et al.* 2005) and are time consuming, the new checks are computationally efficient, making it possible to assess their performance through simulations. We demonstrate the goodness of fit checks in a case study of wetland birds, and assess them in two simulation scenarios with overdispersion in the abundance distribution and in the detection model. The goodness of fit checks are available in an R-package nmixgof.

## 2 Methods

In this section we first introduce the basics of the N-mixture model. In section 2.2 we then develop graphical methods and overdispersion metrics for assessing the fit of N-mixture models. In section 2.3 we demonstrate the use of the goodness of fit checks in a case study on wetland birds in Sweden. Finally, in section 2.4 we investigate the sensitivity of N-mixture models to overdispersion in the abundance and detection models and the ability of the goodness of fit checks to detect violation of the distributional assumptions.

### 2.1 N-mixture models

N-mixture models are a suite of models for abundance data obtained from repeat count surveys at multiple sites (Royle 2004). They model the data as arising from an abundance process describing the spatial variation in the number of individuals among sites and a detection process describing how many of the individuals present at each site are found at each visit. Data come from a set of *R* different sites and for the abundance process it is assumed that the numbers of individuals at sites, *N_i_*, are distributed according to some discrete statistical distribution with probability function *g,*

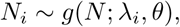

where the draw for each site is independent, *λ_i_* is describing the mean abundance in site *i* which can be a function of covariates, and *θ* is an optional parameter for overdispersion in the abundance distribution. In most applications, *g* is modelled as either a Poisson, a zero-inflated Poisson (ZIP), or as a negative binomial distribution. We will focus on these three mixtures in this paper. For the ZIP mixture we use the parameterisation:

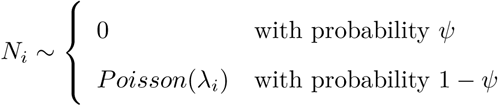

where *ψ* is the probability of an excess zero. For the negative binomial mixture we use the parameterisation:

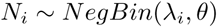

such that the variance of *N_i_* is *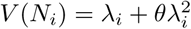*.

For each site observations come in the form of *T* counts, *y_i_*, …, *y*_iT_, and for the detection model it is assumed that the counts are independent binomial draws with population size as index (Royle 2004):

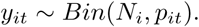

where *p_it_* is the detection probability associated with finding an individual that is present at site *i* at visit *t* and which may vary according to site or visit specific covariates. The design idea underlying this model is that counts are conducted during repeat visits to each site during a period of time for which the local abundance is closed so that at each visit all individuals are present but only a fraction is detected.

Sometimes additional variation in detection is allowed for by letting

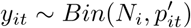

where the *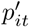* are independently distributed according to a beta distribution

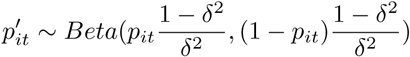

resulting in a beta-binomial detection model. The specific parameterisation in the above equation ensures that *p_it_* is the mean detection probability and that the standard deviation of *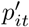* scales linearly with *δ* and is equal to *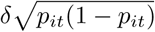*, with 0 *≤ δ ≤* 1.

### 2.2 Checking for over-dispersion and goodness of fit

N-mixture models rely on several crucial assumptions that include population closure within sites at repeat visits (i.e. that the population size *N* remains the same across visits), specific parametric distributions for the detection process and the distribution of abundance as well as functional assumptions about covariate effects.

Checking the fit and assumptions of hierarchical models is difficult in general because distributional and independence assumptions occur at multiple levels in the hierarchy, and through conditioning on unobserved stochastic variables. Current common practice for assessing goodness of fit of N-mixture models, if checked at all, is to use parametric bootstrapping in combination with some goodness of fit statistic, often sums of squares or a Freeman-Tukey statistic (Kéry & Royle 2016). This approach is computationally intensive since in each bootstrap sample the model under investigation needs to be fitted to simulated data a large number of times. In this section we suggest three types of residuals to check the goodness of fit of N-mixture models, as well as two measures of overdispersion relative to a fitted model. The benefit of these over the bootstrap procedure is that i) they are orders of magnitude faster to compute, with computing time measured in terms of seconds rather than hours as is sometimes the case for the parametric bootstrap procedure, and ii) residuals can be used to graphically check a range of assumptions such as overdispersion via quantile-quantile plots (qq plots), residual plots against fitted values to check homoscedasticity, and plots of residuals against covariates to check functional assumptions (Warton *et al.* 2017).

#### 2.2.1 Randomized-quantile residuals

We will define three types of randomized-quantile (rq), or Dunn-Smyth, residuals (Dunn & Smyth 1996). Rq residuals have recently gained popularity in ecological analyses (Warton *et al.* 2016) due to their convenient property that they are normally distributed under the correct model. For sparse count data this means that plots of e.g. residuals against fitted values behave in similar ways to such plots for ordinary linear models which is not the case for standard residuals for count data. That the residuals are indeed normally distributed is also easy to check, for example using qq plots (Warton *et al.* 2016).

The normality of rq residuals is achieved by randomization: For a random count variable *z* with cumulative distribution function (CDF) *F*, they are defined by

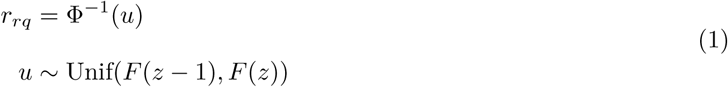

where Φ^−1^ is the inverse of the standard normal CDF and *u* is a value randomly generated from a uniform distribution. To compute rq residuals the function *F* needs to be computed and below we define three variants of rq residuals using CDFs corresponding to different aspects of the data and potentially picking up different aspects of model fit.

##### 2.2.1.1 Marginal rq residuals

For the first type of rq residuals we simply take *F* to be the marginal distribution of the counts (i.e. the distribution of the counts over all possible latent abundances). For the N-mixture model without heterogeneity in *p_it_* and with a Poisson, ZIP or negative binomial mixture distribution, the marginal distribution of each observation comes from the same type of distribution as that used for the abundance mixture. If for example the abundance mixture is *ZIP* (*λ_i_, ψ*), the marginal distribution of each *y_it_* is *ZIP* (*p_it_λ_i_, ψ*). In these cases the randomized-quantile residuals can be easily computed using the definition above (eq. 1).

For beta-binomial detection models the marginal distribution is to our knowledge not available in closed form but can be computed by numeric summation over *N* using

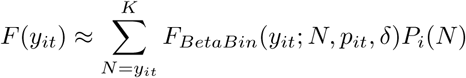

where *K* is large enough that the contribution from larger *N* can be ignored, *F_BetaBin_* is the CDF of the beta-binomial, and *P_i_*(*N*) is the probability that the abundance at site *i* is equal to *N* given by the abundance distribution. This is similar to how the likelihood of the N-mixture model can be approximated by a finite sum (Royle 2004).

A property of the marginal rq residuals computed from an N-mixture model is that residuals from the same site are not independent because the counts are not. Hence they should not be used directly in qq plots which assume independent observations. However sets of residuals containing only one residual from each site are independent in the same way that sets of counts are, and separate qq plots can be drawn for each set. Since there is one marginal rq residual per observation, they can be plotted against visit specific detection covariates as well as against site specific covariates.

##### 2.2.1.2 Site-sum rq residuals

The second type of residuals we propose is defined from the marginal distribution of the sum of the counts within each site *y_Si_* =Σ *_t_y_it_*. The marginal CDF for the site sums can be computed numerically using

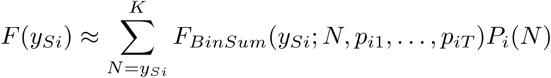

where *F_BinSum_* is the CDF of a sum of independent binomial variables, all with the same index *N* but potentially different probabilities *p_it_*. If the *p_it_* are all the same *F_BinSum_* is simply the cumulative probability function of a binomial distribution with index *T N* but if the *p_it_* are not all identical then *F_BinSum_* is more complex. In the general case it can be computed by brute force as a numeric sum:

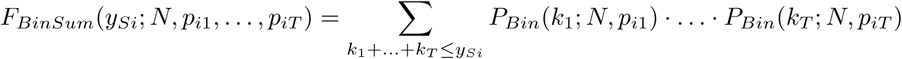

where *P_Bin_* is the probability function of the Binomial distribution. The same computation may be used for beta-binomial detection models by replacing *P_Bin_* with *P_BetaBin_*.

The idea of aggregating counts across sites is to make the residuals independent and to potentially increase their informativeness in cases where counts are sparse. Since there is one site-sum residual per site, they can be used in plots against site-specific covariates.

##### 2.2.1.3 Observation rq residuals

We also explored a third type of residuals that we refer to as observation residuals. The idea is to compute residuals from the observation model only by conditioning on the abundances, with the intent of more specifically checking the detection part of the model. Since the abundances are not directly available from a fitted model we use a random sample of abundances from the empirical Bayes distribution (the distribution of the abundances given the data and under the parameters obtained by maximum likelihood) for the conditioning. That is, residuals were computed using the binomial or beta-binomial CDF with *N_i_* equal to a draw from the empirical Bayes distribution. The random draw introduces additional stochasticity to the residuals which is likely to reduce their power to some degree.

#### 2.2.2 Measures of overdispersion

The parametric bootstrap procedure used to check goodness of fit mentioned above has also been used to provide a measure of overdispersion (Kéry & Royle 2016) through

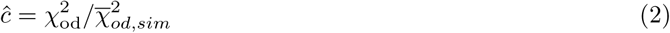

where *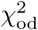* is a goodness of fit statistic computed from a model fit to the data and *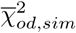* is the mean of the same statistic computed from fits of the model to data simulated from the model using parameters estimated from the original data. Under the correct model the expectation of *ĉ* is 1 while we would expect *ĉ* to be greater than 1 if the data are over-dispersed relative to the fitted model (and less than 1 if they are under-dispersed). Clearly this is a computationally expensive calculation and thereby difficult to evaluate through simulations. The goal in this section is to find similar measures with less of a computational burden, and whose behaviour we will explore in simulations in a later section.

For measures of discrepancy between the observed data and a fitted model we use chi-square type statistics based on Pearson residuals which have the form (Hilbe 2011):

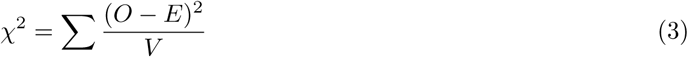

where *V* is the variance of the observations *O* and *E* is its expectation. The statistic differs from the standard chi-square statistic which has the form

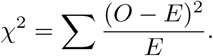

The former collapses to the latter when the variance is equal to the mean, such as when *O* are counts from a Poisson distribution with mean *E*. The statistic based on Pearson residuals has the advantage that the expectation of the terms in the sum are 1 under the correct model which is not the case for the standard chi-square statistic in general (e.g. under a negative binomial model). We will use this feature here to define overdispersion metrics that have mean 1 under the correct model. We will consider two variants of overdispersion measures, one based on marginal Pearson residuals and the other based on site-sum Pearson residuals.

##### 2.2.2.1 Marginal *ĉ*

For the marginal measure of overdispersion we use the chi-square statistic based on Pearson residuals (eq. 3) computed over each observation:

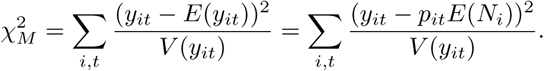

The general expression for the variance of the counts with beta-binomial detection is

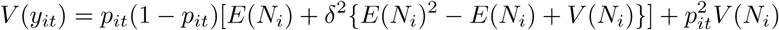

where *E*(*N_i_*) and *V* (*N_i_*) are the mean and variance given by the abundance mixture (a derivation of this formula is given in Appendix 1). For the simplest case with Poisson distributed abundances (*E*(*N_i_*) = *V* (*N_i_*) = *λ_i_*) and binomial detection (*δ* = 0) the variance reduces to

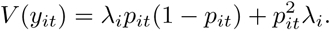

From *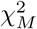* we define the marginal overdispersion measure as

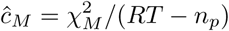

where *n_p_* is the number of parameters of the model and *RT* is the product of the number of sites and the number of visits, i.e. the total number of counts.

##### 2.2.2.2 Site-sum *ĉ*

We define t he s ite-sum m easure o f overdispersion by computing t he chi s quare s tatistic (eq. 3) f or Pearson residuals of the summed counts across sites:

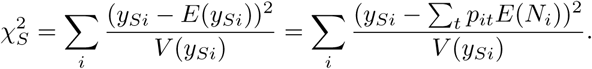

The variance of the summed counts in the above equation is

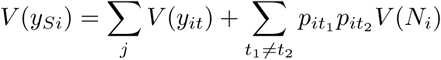

From this we define t he s ite-sum m easure o f overdispersion by a gain d ividing by t he number o f t erms i n the sum (*R*) less the number of parameters (*n_p_*):

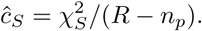

### 2.3 Case study: Northern shoveler

To illustrate the performance of the residuals and overdispersion metrics above, we analyse data from a wetland survey conducted in May and June of 2016 at 50 wetland sites across southern Sweden. Most sites (90%) were visited 10 times during a three week period, split between 5 visits by each of two observers, but some sites had fewer visits. The number of individuals for each of 70 bird species associated with wetlands was recorded on each visit. Here, we use counts for Northern shoveler (Fig. S1), a dabbling duck moderately common in lakes and wetlands in southern Sweden. We fit six N-mixture models to the data using combinations of Poisson (P), ZIP and negative binomial (NB) abundance mixtures and binomial (B) and beta-binomial (BB) detection. Hereafter the models will sometimes be referred to using abbreviations such as BB-ZIP with prefix denoting the detection model and suffix denoting the abundance distribution. All models included two covariates for abundance, the log transformed total area of water at the wetland representing its size and the latitude of the wetland, and two covariates for detection, the date of the survey and the percentage of reed cover at the wetland as a proxy for visibility. All covariates were introduced as linear functions on the log (abundance) and logit scale (detection) and were standardized to mean 0 and standard deviation 1 prior to analyses. We fitted models with binomial detection using the R-package unmarked (Fiske & Chandler 2011) and models with beta-binomial detection using custom code.

The N-mixture model as implemented in unmarked approximates the likelihood by truncating an infinite sum over all possible values of *N*. The upper bound, *K*, needs to be set when fitting the model, but it is known that estimates can be unstable to changes in this bound, possibly due to maximum likelihood estimates of abundance being infinite (Dennis *et al.* 2015). We used a numeric upper bound *K* = 400 for abundance in the calculation of the likelihoods but also fitted the same models a second time using *K* = 1000 to check if the estimates were stable to this numeric cutoff.

#### 2.3.1 Results of case study

Estimates under the Poisson and ZIP abundance mixtures were not sensitive to the numerical cutoff *K* while this was the case for both models with an NB mixture. The estimates obtained for the NB mixtures are thus not maximum likelihood estimates, and estimates of abundance will increase and those of detection decrease as *K* is increased. We will refer to them as truncated estimates. Models with binomial and beta-binomial detection give similar estimates under the same abundance mixture but the estimates differ among abundance mixtures (Fig. 1).

**Figure 1:**
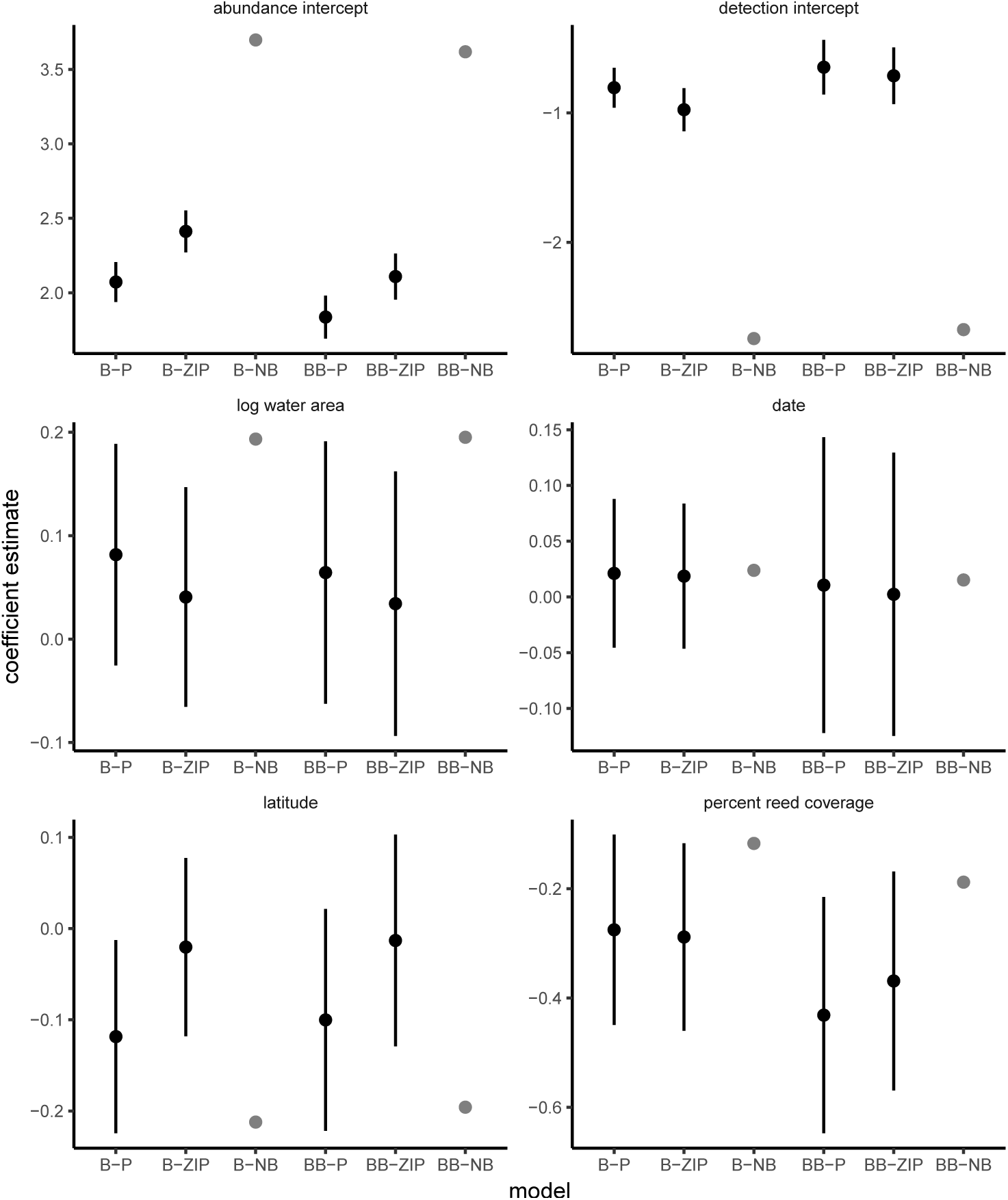
Estimates and 95% confidence intervals for intercepts and covariates coefficients for abundance (left panels) and detection (right panels) of the models fitted to Northern shoveler data. Prefix B and BB refers to, respectively binomial and beta-binomial detection models. Suffix P, ZIP and NB refers to Poisson, zero-inflated Poisson, and negative binomial abundance mixtures. Estimates under the NB mixtures are unstable and not maximum likelihood estimates. Truncated point estimates are given in gray for K=400 for those models, but confidence intervals are omitted.

Qq plots of site-sum randomized quantile residuals show that models with Poisson or ZIP mixtures provide poor fits to the data since the quantiles deviate clearly from the identity line (Fig. 2), while the truncated estimates of the NB mixtures appear adequate (Fig. 2). The qq plots for the Poisson mixtures indicate that the largest residuals are larger and the smallest smaller than would be expected under Poisson mixtures while the qq plots for the ZIP mixtures show some improvement in terms of explaining the smallest (zero) observations, but is still at loss in explaining larger counts. Similar patterns are seen for the marginal rq residuals (Fig. S2). The *ĉ* measures similarly indicate substantial overdispersion (*ĉ >>* 1) for the Poisson and ZIP mixtures but not for the truncated NB estimates (Table 1). Overdispersion is stronger according to *ĉ_M_* than *ĉ_S_* (Table 1). Similarly, AIC values indicate a poor fit of the Poisson and ZIP mixtures relative to the truncated NB mixture estimates (Table 1). AIC in addition suggest a poor fit of the truncated B-NB model relative to the truncated BB-NB model which is not picked up by the qq plots of site-sum residuals or *ĉ_S_*. Qq plots of observation residuals however do suggest lack of fit of the truncated B-NB model (Fig. 3). Qq plots of observation residuals for the truncated BB-NB model show no obvious lack of fit (Fig. 3).

**Table 1:** Estimates of overdispersion for fits to Northern shoveler data.

| model | $\hat{c}_S$ | $\hat{c}_M$ | AIC |
| --- | --- | --- | --- |
| B-P | 11.0 | 5.3 | 2026.3 |
| B-ZIP | 4.4 | 3.3 | 1915.6 |
| B-NB | 0.9 | 1.3 | 1601.6 |
| BB-P | 9.2 | 2.9 | 1789.5 |
| BB-ZIP | 4.6 | 2.2 | 1719.8 |
| BB-NB | 0.9 | 0.8 | 1568.3 |

**Figure 2:**
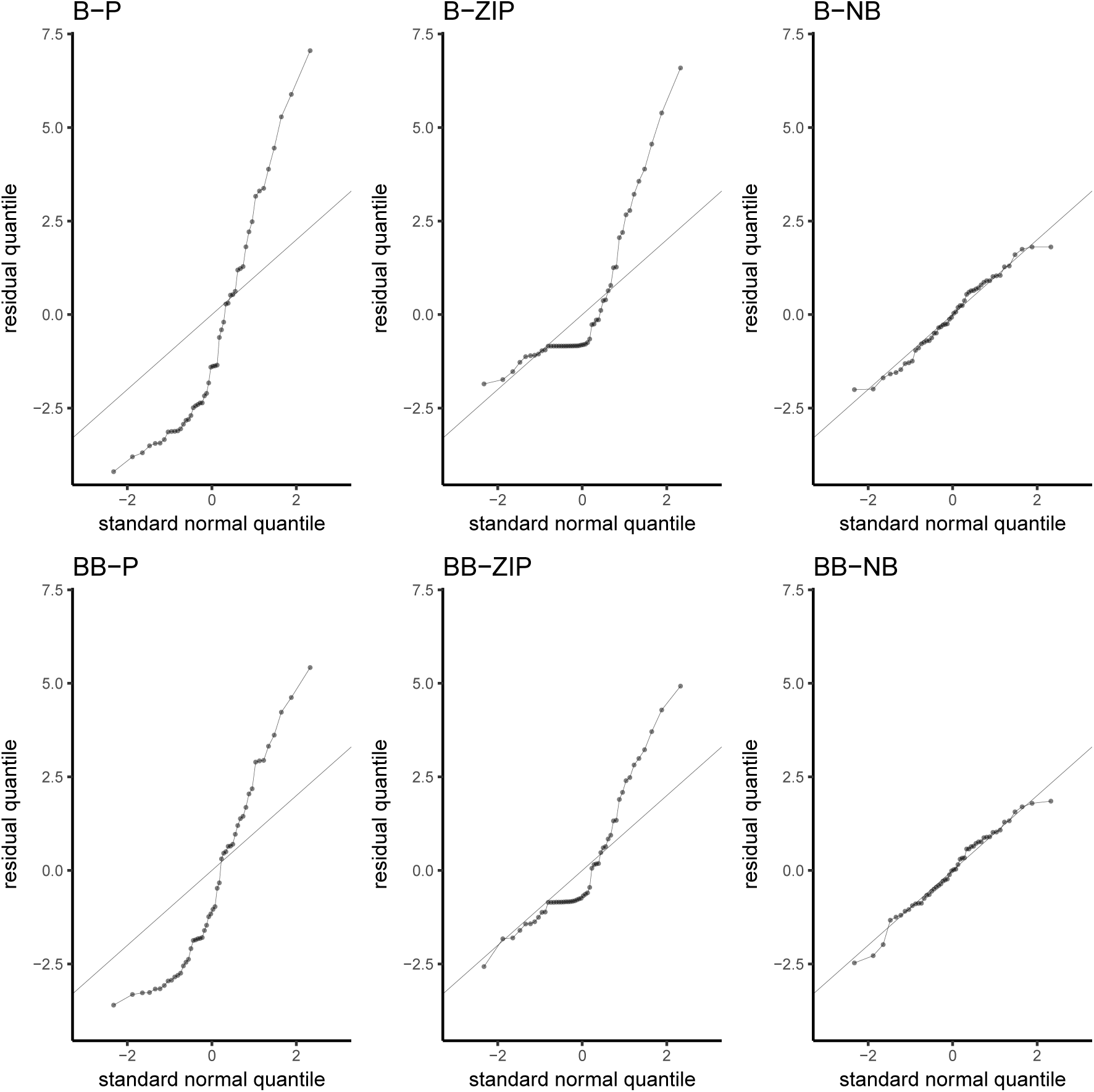
QQ plots of site-sum randomized-quantile residuals against standard normal residuals for fits of models to the Northern shoveler data. Under a good fit residuals should be close to the identity line (gray). Prefix B and BB refers to, respectively binomial and beta-binomial detection models. Suffix P, ZIP and NB refers to Poisson, zero-inflated Poisson, and negative binomial abundance mixtures.

**Figure 3:**
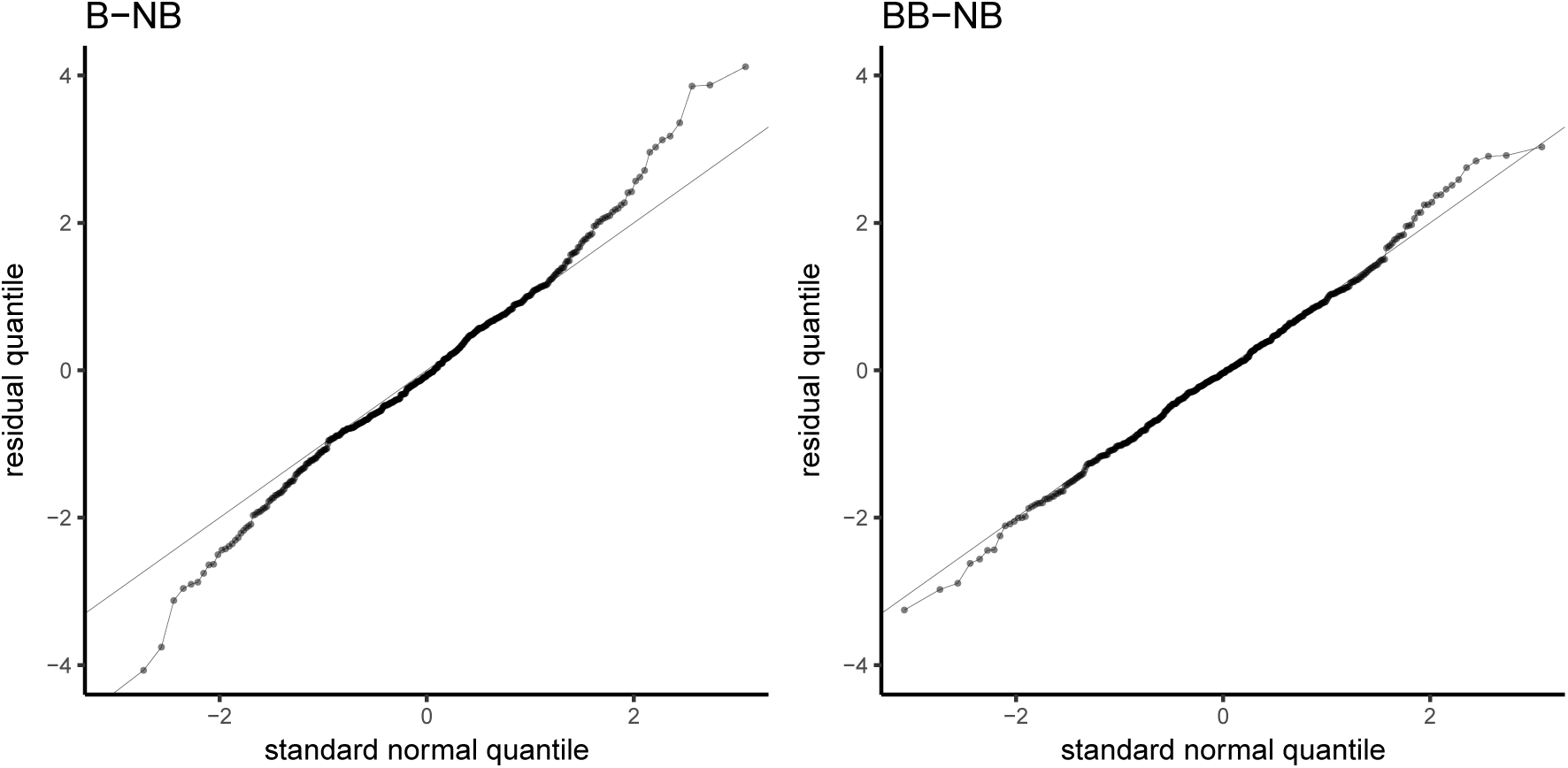
QQ plots of observation randomized quantile residuals against standard normal residuals for fits of binomial and beta-binomial NB models to the Northern shoveler data. Under a good fit residuals should be close to the identity line (gray). B and BB refers to, respectively binomial and beta-binomial detection models, while NB refers to the negative binomial abundance mixture.

These results leave us in a quandary. The NB mixtures give unstable estimates and cannot be used for inferences about abundance, and the poor fit of the Poisson and ZIP mixtures suggest that we cannot use estimates from these models for reliable inference either. To check if the reason for the poor fit of the Poisson and ZIP mixtures might be due to incorrect functional covariate relationships we plot rq residuals against each of the covariates for the BB-ZIP model, which has the best fit among the models with stable estimates (Fig. S3). Since there is no clear pattern in the residuals as a function of covariates for this model there appears to be no simple correction to improve its fit. The conclusion from this case study therefore has to be that we are not able to find an adequately fitting N-mixture model that provides reasonable estimates for the data at hand. The seemingly decent fit using the truncated estimates from the NB mixtures on the other hand suggest that an analysis of relative abundances with generalized linear mixed models accounting for overdispersion could be fruitful (Barker *et al.* 2017), but we do not pursue this further here.

### 2.4 Simulations

To investigate the properties of our goodness of fit checks, and how they relate to potential bias in parameter estimates, we ran two simulation scenarios, one where there is overdispersion in the abundance distribution relative to the Poisson distribution and one where there is overdispersion in detection relative to the binomial distribution such that detection probabilities vary independently among sites and visits.

#### 2.4.1 Scenario 1: Overdispersed abundance

We simulated data over 200 sites, each visited 5 times, using a binomial detection model with *p_it_* set to 0.25 for all visits and sites and with a constant expected abundance across all sites *λ_i_* = 10. To investigate effects of overdispersion we used a negative binomial abundance distribution and varied the overdispersion coefficient *θ* from 0 to 2 in steps of 0.25. Thus, data were generated using a B-NB model. For each value of *θ* 500 data sets were generated. For each simulated data set we fit a B-P, B-ZIP, B-NB (which in this simulation is the correct model), and a BB-P N-mixture model, each with a single intercept for detection and abundance but no covariates. The models with binomial detection (B-P, B-ZIP, and B-NB) were fitted in unmarked while the BB-P model was fitted using custom R-code.

In addition, we fitted a second set of models that were identical to the ones described above except for the addition of a single covariate for abundance. The covariate was generated from a standard normal distribution and was used in the fitted models but was unrelated to the simulated data. These three models with covariates were fitted in order to investigate if overdispersion might lead to finding spurious effects of covariates (Richards 2008).

We used a numeric cutoff value *K* = 200 for the calculation of the likelihood during model fitting. To check for stability of estimates with respect to *K* we additionally fitted each model using a *K* value of 400 and classified estimates as stable if the abundance intercept between the two *K* values differed by less than 0.01.

For all the fitted models we retrieved parameter estimates, AIC, and also computed *ĉ_M_* and *ĉ_S_*. As a rough estimate of the power of the qq plots to detect non-normality in the randomized quantile residuals we computed the p-value from a Shapiro-Wilks test of normality for the site-sum and observation residuals (this was not done for the marginal residuals because they are not independent among visits). We do not recommend this procedure in applications but used it here to obtain a crude but objective measure of power of the residuals to detect lack of fit. In applications we suggest using graphical checks via qq plots and plots of residuals against fitted values and covariates because such checks provide more information about the nature of the lack of fit than a p-value does.

#### 2.4.2 Scenario 2: Overdispersed detection

In the second scenario we explored the effects of overdispersion in detection relative to the binomial distribution. The setup in this scenario is similar to the setup in scenario 1, except that we used a Poisson abundance mixture and a beta-binomial detection model to simulate data (i.e. a BB-P model). We varied *δ*, i.e. the amount of variation in the detection probability, from 0 to 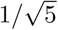. The upper bound was chosen so that the distribution of the detection probability has an interior mode for all values of *δ* except for *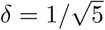* where the mode is at 0. We fitted the same models as in scenario 1.

#### 2.4.3 Simulation results: scenario 1

Nearly all model fits converged and were stable with respect to *K* in this scenario (Fig. 4a). As expected, fitting the true B-NB model provided the least bias, nearly nominal confidence interval coverage for the covariate effect, *ĉ* measures close to 1, and rejected the normality test for the rq residuals in proportion to the alpha level (Fig. 4).

**Figure 4:**
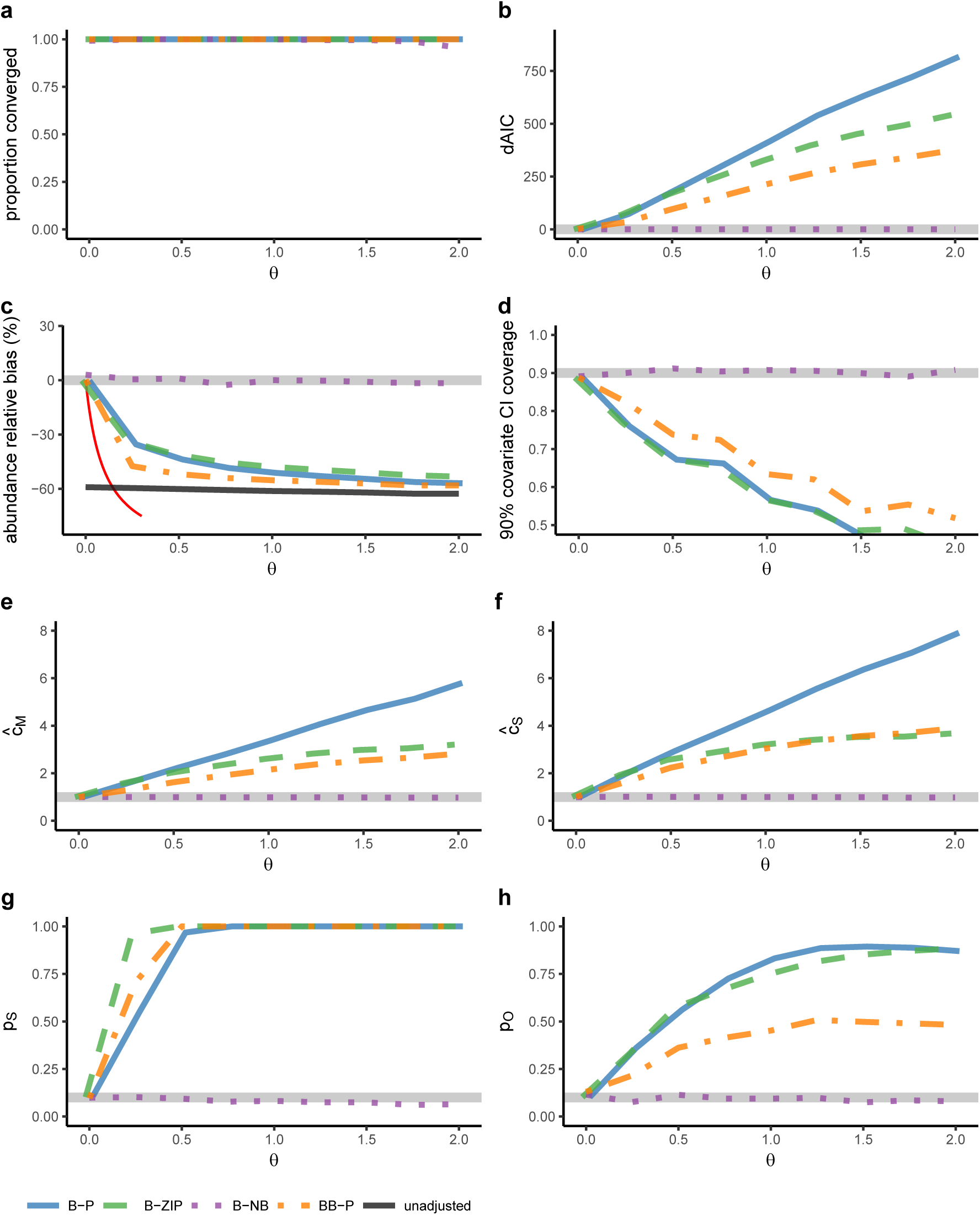
Results for binomial Poisson (B-P, blue), binomial ZIP (B-ZIP, green), binomial NB (B-NB, magenta), and beta-binomial Poisson (BB-P, orange) models fitted to data simulated from a negative binomial mixture with binomial detection (scenario 1) as a function of the overdispersion *θ*. Grey lines give the reference level in each panel. a) Proportion of simulations for which estimates where stable relative to the numerical cutoff K and for which the optimization routine converged. b) Average difference in AIC between each model and the fitted correct B-NB model. c) Relative bias in estimated mean abundance. Black line gives estimates not adjusted for imperfect detection, computed as the mean of the maximum counts at each site. The red line gives the theoretical bias of the BB-P model by matching moments. d) Proportion of Wald confidence intervals (90%) for the covariate effect that cover the true value (0). e) Marginal overdispersion measure. f) Site-sum overdispersion measure. g) Proportion of simulations for which a normality test (Shapiro) computed from site-sum rq residuals was rejected at the 10% level. h) Proportion of simulations for which a normality test (Shapiro) computed observation rq residuals was17rejected at the 10% level.

The B-P, B-ZIP and BB-P models strongly underestimated abundance for high levels of overdispersion with a relative bias of less than -50% for the B-P, B-ZIP and BB-P models (Fig. 4c). The strongest bias was given by the BB-P model. These levels of bias are of similar magnitude to estimates not adjusted for detection, which had a relative bias of around -60%. Overdispersion also led to poor confidence interval coverage for the spurious covariate effect, except when fitting the correct model (Fig. 4d).

Lack of fit relative to the true B-NB model was readily identified by AIC in the simulations (Fig. 4b). Absolute lack of fit was similarly well identified by *ĉ_M_* and *ĉ_S_* but the latter estimates of overdispersion were higher (Fig. 4e and f). Considerable bias in the abundance estimates (more than 30%) was associated with average *ĉ_M_* and *ĉ_S_* as low as 1.5.

Normality tests of the site-sum rq residuals rejected incorrect models at high rates (Fig. 4g), but observation rq residuals had considerably lower power (Fig. 4h).

#### 2.4.4 Simulation results: scenario 2

Most model fits in scenario 2 converged and were stable with respect to *K*, except under the B-NB model that failed for almost all simulated data sets when *δ >* 0.2 (Fig. 5a). Properties of the model fits like bias, coverage etc. were computed only from fits that converged and were stable with respect to *K*.

**Figure 5:**
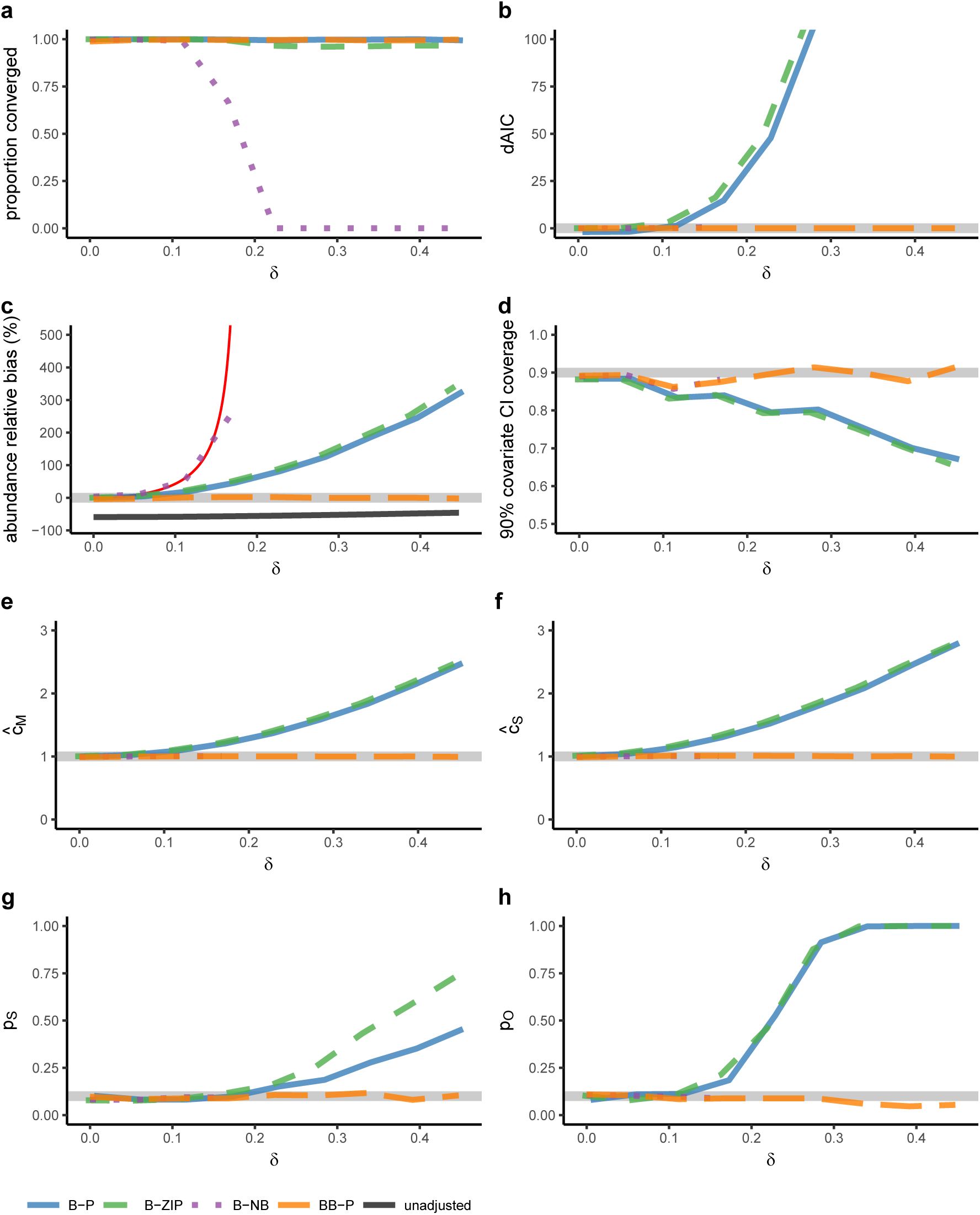
Results for binomial Poisson (B-P, blue), binomial ZIP (B-ZIP, green), binomial NB (B-NB, magenta), and beta-binomial Poisson (BB-P, orange) models fitted to data simulated from a Poisson mixture with beta-binomial detection (scenario 2) as a function of the amount of variation in detection probability *δ*. Grey lines give the reference level in each panel. a) Proportion of simulations for which estimates where stable relative to the numerical cutoff K and for which the optimization routine converged. b) Average difference in AIC between each model and the fitted true BB-P model. c) Relative bias in estimated mean abundance. Black line gives estimates not adjusted for imperfect detection, computed as the mean of the maximum counts at each site. The red line gives the theoretical bias of the B-NB model by matching moments. d) Proportion of Wald confidence intervals (90%) for the covariate effect that cover the true value (0). e) Marginal overdispersion measure. f) Site-sum overdispersion measure. g) Proportion of simulations for which a normality test (Shapiro) computed from site-sum rq residuals was rejected at the 10% level. h) Proportion of simulations for which a normality test (Shapiro)computed observation rq residuals was rejected at the 10% level.

The B-NB model, when it converged, strongly overestimated abundance even for small amounts of variation in the detection probability, while the B-P and B-ZIP models strongly overestimated abundance when the variation in detection probability was larger (Fig. 5c). The correct beta-binomial Poisson model (BB-P) provided unbiased estimates. Confidence intervals for the spurious covariate had acceptable coverage for moderate variation in the detection probability but declined as that variability increased except under the correct model (Fig. 5d).

The overdispersion measures *ĉ_M_* and *ĉ_S_* performed similarly in detecting lack of fit. They were unable to indicate lack of fit of the strongly biased B-NB model but did increase for the B-P and B-ZIP models as the variation in the detection probability increased (Fig. 5e and f). However, even when abundance was estimated at twice its true value (100% relative bias) under these models, the overdispersion measures were only around 1.5. These metrics therefore struggled to indicate lack of fit, and overdispersion metrics only slightly larger than 1 could correspond to very strong bias in estimated abundance.

Normality tests of rq residuals similarly failed to detect lack of fit for small to moderate variation in the detection probability. For large variation in the detection probability the test of the observation rq residuals did often detect lack of fit and had better power than the test of the marginal rq residuals (Fig. 5g and h).

AIC had better performance in determining relative lack of fit of the B-P and B-ZIP model in relation to the true BB-P model, but was unable to distinguish between the B-NB model and the true model (Fig 5b).

#### 2.4.5 Approximating the BB-P N-mixture model with a B-NB model

The inability of the overdispersion measures to diagnose lack of fit of the B-NB model in scenario 2, the small difference in AIC between this model and the true BB-P model for moderate values of *δ*, and the collapse at large values of *δ*, can be understood through approximating the BB-P model with a B-NB model. Barker *et al.* (2017) recently used moment matching to show that Poisson and negative binomial N-mixture models with a binomial detection model can be approximated by a double Poisson regression model, the latter lacking any notion of a latent abundance. Using moment matching, we show in Appendix 1 that an N-mixture model with beta-binomial detection and a Poisson abundance mixture can be approximated by another N-mixture model with binomial detection and a negative binomial abundance mixture where the abundance is inflated as long as *δ*^2^ *< p/*(*λ - λ p*). In other words, data from a BB-P model will look identical to data from a B-NB model with higher abundance in terms of means, variances and covariances for such values. Because of this it is difficult to distinguish between overdispersion in the detection probability and overdispersion in abundance. The only chance to separate between them is therefore to resort to more subtle properties of the models given by their higher order moments.

This explains why the overdispersion measures *ĉ_M_* and *ĉ_S_* cannot detect lack of fit in scenario 2 since they only depend on the first and second order moments of the models. It also gives a justification for the breakdown of the B-NB model around *δ* = 0.2 in Fig. 5. The moment matching gives negative *p* for the B-NB model if *δ >* 0.18. For these values of *δ*, the best moment approximation is therefore *p* = 0 and *λ* = *∞*. For values of *δ <* 0.18 the expected bias from the B-NB moment approximation matches the bias in the simulations (Fig. 5c).

The above approximation also suggests that the BB-P model could underestimate abundance and provide a decent fit to data that are generated from a B-NB model with the same moments as long as *δ*^2^ *< p/*(*λ - λp*). For larger values of *δ* there is no matching B-NB model but we show in Appendix 1 that for such *δ* there is a range of BB-NB N-mixture models with the exact same moments as the BB-P model. This range contains one model for each possible value of abundance larger than *λ*. Hence, data that have first and second moments that matches the BB-P model could have been generated from a model with overdispersion in both abundance and detection with a much higher abundance than the BB-P model would suggest.

## 3 Discussion

N-mixture models provide an appealing framework for learning about absolute rather than relative abundance of populations from count data alone, but this comes at the price of a very strong reliance on model assumptions. Count data by themselves contain only minimal information about absolute abundances (Knape & Korner-Nievergelt 2015; Barker *et al.* 2017) and our results, and some results of previous studies (Martin *et al.* 2011; Toribio *et al.* 2012), show that this leads to N-mixture models often being sensitive to even small amounts of model mis-specification. As a result, estimates of abundance and detection can be severely biased and inference about effects of covariates misleading if model assumptions are not met to a satisfactory degree. In light of this, finding a model that adequately fits the data is necessary for reliable inferences about abundance using N-mixture models. The diagnostic tools proposed here are designed to evaluate the goodness of fit of N-mixture models.

Our results show sensitivity of estimated abundances to overdispersion in the abundance mixture and, as previously shown (Martin *et al.* 2011), in the detection probability if the overdispersion is not accounted for. Not accounting for overdispersion in the abundance mixture leads to underestimating actual abundance while not accounting for random variation in the detection probability leads to overestimating abundance. In our simulations, site-sum rq residuals and marginal and site-sum overdispersion measures were effective in detecting lack of fit caused by overdispersion in the abundance mixture. However, average values of the overdispersion metrics as small as 2 or less corresponded to underestimating abundance by 30% on average. We found detecting lack of fit due to overdispersion in the detection probability to be more challenging. Lack of fit of a binomial detection model due to random variation in the detection probability among sites and visits was only reliably detected at levels of overdispersion where bias was already large. Rq residuals and overdispersion metrics had no power to detect lack of fit of the negative binomial model even when abundance was overestimated by over 300%, but had some power to detect lack of fit of the binomial Poisson and ZIP models for high variability in the detection probability. Like for lack of fit due to overdispersion in abundance, small values of the overdispersion metrics can correspond to strong bias in estimated abundance.

Problems with detecting lack of fit due to variation in the detection probability occur despite the fact that we used a large sample size of 200 sites and 5 repeat visits in our simulation, and are not simply due to a poor choice of goodness of fit metrics. The problems arise due to a fundamental similarity between alternative model structures for the same data leading to difficulties in distinguishing between models. We show in Appendix 1 that the first and second order moments of the negative binomial N-mixture model can be matched exactly to the moments of a beta-binomial Poisson N-mixture model for small to intermediate variability in the probability of detection. This correspondence explains why detecting lack of fit is problematic for this model since higher order moments are needed to separate between them. That is, data from a negative binomial model and a beta-binomial Poisson model can behave in much the same way and are therefore difficult to separate. While it is possible that alternative goodness of fit metrics that are more efficient in detecting lack of fit due to variation in the detection probability could be designed, this will be a hard and sometimes impossible problem to solve, especially for limited sample sizes such as a low number of repeat visits.

Barker *et al.* (2017) recently used moment matching to show that alternative data generating mechanisms can give rise to data that are similar to the binomial Poisson and negative binomial N-mixture models. The moment matching here extends these results to beta-binomial models, and does so within the extended framework of beta-binomial negative binomial N-mixture models to show that a wide range of different abundances can give rise to similar data. This is concerning for the robustness of estimates of abundance using the beta-binomial model. Most real data sets would be expected to contain overdispersion (or sometimes underdispersion) in both the detection and the abundance process. The beta-binomial negative binomial N-mixture model provides one framework for such data, but we have shown that this framework is identifiable only by considering moments of the models beyond those of the second order (means, variances and covariances) so that resorting to arguably subtle properties of the models would be required to identify abundance.

The bias of the N-mixture model under mis-specification depends on parameter values. We used a moderately low detection probability (*p* = 0.25) and a high abundance (*λ* = 10) in our simulations. The moment matching suggests that if the detection probability is higher or abundances lower, the biases will be smaller and the N-mixture model more robust. The problem in practice is that these quantities are unknown. It seems tempting to rely on estimated detection probabilities and abundances from a fitted model to determine that one is in the parameter region where estimates are robust, but it is clear from the simulations that such an approach is not reliable. In scenario 1, estimated detection probabilities under models ignoring overdispersion in abundance were much higher than the detection probabilities used to simulate the data. Our suggestion is to instead fit multiple N-mixture models with and without overdispersion to the same data. In the parameter region where the N-mixture model is more robust, the different models are expected to provide similar although not necessarily identical estimates. In cases where the different models give similar abundances and fit the data well, the estimation issues discussed here may therefore be less of a problem.

Variability in the detection probability led to failure of the negative binomial N-mixture model such that it provided practically infinite estimates of abundance as this variability increased. This happened in our simulations when matching the moments of the negative binomial N-mixture model to the beta-binomial model suggests a negative probability. Thus our results give a mechanism through which the negative binomial model can fail to provide finite estimates of abundance, a problem that has been commonly observed in case studies and in simulations (Dennis *et al.* 2015; Kéry & Royle 2016).

The goodness of fit checks discussed here for binomial N-mixture models are easily extended to multinomial N-mixture models (Kéry & Royle 2016). Site-sum rq residuals and overdispersion metrics may for example be defined for the sum of counts over all the observed categories of the multinomial. In distance sampling this equates to the total number of individuals detected across all distances at each site, and our *ĉ_S_* measure defined in this way would correspond to the *ĉ* metric for distance sampling given by Johnson *et al.* (2010) in the case of Poisson distributed abundances.

### 3.1 Conclusions

Some studies have questioned the utility of the N-mixture framework (Hunt *et al.* 2012; Couturier *et al.* 2013; Barker *et al.* 2017). Our results extend concerns about robustness to N-mixture models with beta-binomial detections, which have been argued to be more robust than their binomial counterparts (Martin *et al.* 2011). We agree with Barker *et al.* (2017) that count data lacking additional information about detection probabilities are often better treated as indices of relative abundance than used to estimate absolute abundance. By treating data as indices one can get around the instabilities often associated with the N-mixture model and utilize more standard frameworks like the generalized linear or additive mixed models (Link & Sauer 1997; Fewster *et al.* 2000; Knape 2016) with their suite of methods for assessing model fit (Barker *et al.* 2017). Alternatively, detection probabilities in the binomial N-mixture model may be calibrated using additional information about detections for some or all sites, e.g. through removal (Farnsworth *et al.* 2002) or distance sampling (Johnson *et al.* 2010) protocols. If one despite the concerns with robustness uses binomial N-mixture models for estimating absolute abundance one should make sure that the final model provides a good fit to the data. Doing so will provide some steps towards reducing the risk of strongly biased estimates. Our goodness of fit checks can be used to this end and are available in an R-package nmixgof compatible with unmarked.

## Acknowledgements

We thank Marc Kéry for providing valuable comments that improved the paper. JK was funded by grant 621-2012-4076 from the Swedish Research Council VR and DA by grant 942-2015-287 from FORMAS. The wetland survey was funded by a grant from FORMAS to TP. FB was supported by LabEx COTE (ANR-10-LABX-45).

